# *Wolbachia* incompatible insect technique program optimization over large spatial scales using a process-based model of mosquito metapopulation dynamics

**DOI:** 10.1101/2024.07.01.601531

**Authors:** Preston Lim, Alex R Cook, Somya Bansal, Jo Yi Chow, Jue Tao Lim

## Abstract

**Background:** *Wolbachia* incompatible insect technique (IIT) programs have been shown in field trials to be highly effective in suppressing populations of mosquitoes that carry diseases such as dengue, chikungunya and Zika. However, the frequent and repeated release of *Wolbachia*-infected male mosquitoes makes such programs resource-intensive. While the need for optimization is recognized, potential strategies to optimize releases and reduce resource utilization have not been fully explored.

**Methods:** We developed a process-based model to study the spatial-temporal metapopulation dynamics of mosquitoes in a *Wolbachia* IIT program, which explicitly incorporates climatic influence in mosquito life-history traits. We then used the model to simulate various scale-down and redistribution strategies to optimize the existing program in Singapore. Specifically, the model was used to study the trade-offs between the intervention efficacy outcomes and resource requirements of various release program strategies, such as the total number of release events and the number of mosquitoes released.

**Results:** We found that scaling down releases in existing sites from twice a week to only once a week yielded minimal changes in suppression efficacy (from 91% to 85%), while requiring 45% fewer mosquitoes and release events. Additionally, redistributing mosquitoes from already suppressed areas and releasing them in new areas once a week led to a greater total suppressive efficacy (88% compared to 65%) while also yielding a 16% and 14% reduction in the number of mosquitoes and release events required respectively.

**Conclusions:** Both scale-down and redistribution strategies can be implemented to significantly reduce program resource requirements without compromising the suppressive efficacy of IIT. These findings will inform planners on ways to optimize existing and future IIT programs, potentially allowing for the wider adoption of this method for mosquito-borne disease control.

## Background

Mosquitoes are vectors for several tropical diseases, including malaria, dengue, chikungunya, Zika, and lymphatic filariasis. It is estimated that up to 700 million people worldwide are infected and more than a million people die from mosquito-borne diseases every year (World Health Organization: WHO, 2020). At present, vector control methods such as breeding site reduction and insecticide spraying are the most effective tools for combating these mosquito-borne diseases, as pharmaceutical interventions are not widely available (World Health Organization: WHO, 2014). Given growing concerns of emerging insecticide resistance (World Health Organization: WHO, 2014), public health organizations and governments worldwide have been exploring novel vector control methods to supplement existing toolkits.

One promising approach for some diseases is the use of mosquitoes infected with *Wolbachia*, a naturally occurring intracellular alpha-proteobacterium that is commonly found in insect species (Bourtiz et al., 2014). *Wolbachia* has been found to confer the mosquito host with inhibitory effects against certain viral pathogens, thereby reducing the transmission potential for dengue, chikungunya, and Zika (Moreira et al., 2009; Aliota et al., 2016). Population replacement trials, in which both male and female *Wolbachia*-infected mosquitoes are released, have also been conducted in Australia, Vietnam, Malaysia and Indonesia, resulting in significant decreases in dengue incidence (Ant et al., 2022).

*Wolbachia* also affects arthropod reproduction by inducing cytoplasmic incompatibility. When male insects infected with *Wolbachia* mate with females that either lack *Wolbachia* or have different *Wolbachia* strains, cytoplasmic incompatibility can cause the offspring to die during the early stages of embryonic development (Engelstädter & Telschow, 2009). *Wolbachia-*mediated cytoplasmic incompatibility has been tested in field trials in China, the USA, and Singapore through the incompatible insect technique (IIT), where only male mosquitoes with the *Wolbachia* infection are released, and has been found to effectively suppress the mosquito population (Zeng et al., 2022; Crawford et al., 2020; The Project Wolbachia - Singapore Consortium & Ng, 2021; Bansal et al., 2023) and consequently reduce dengue incidence (Lim et al., 2023; Lim et al., 2024). A cluster-randomized controlled trial is also underway in Singapore to provide gold-standard evidence of the technology’s epidemiological efficacy (Ong et al., 2022). IIT is also sometimes supplemented with the sterile insect technique, in which a low-dose of sterilizing radiation is applied to counteract the accidental release of fertile *Wolbachia*-infected female mosquitoes due to imperfect sex-sorting methods (Ong et al., 2022).

Although cost-effective in specific settings (Soh et al., 2021), *Wolbachia*-based IIT programs are resource intensive since they require the repeated and frequent release of large quantities of male *Wolbachia*-infected mosquitoes over the treatment period (Matsufuji & Seirin-Lee, 2023). This is especially in cases where high-fidelity sex-sorting techniques are employed to ensure a low risk of accidentally releasing female mosquitoes (Crawford et al., 2020). In Singapore, for example, a total of 11 million male IIT mosquitoes are released every week (Ganesan, 2024), with each treatment site receiving mosquitoes twice a week and some sites requiring releases at different floors in high-rise apartment blocks (Ong et al., 2022). As such, identifying optimization strategies to safely scale down the IIT release program once the wild-type mosquito population is effectively suppressed is crucial. Scale-down strategies can also reduce the risk of releasing female *Wolbachia* mosquitoes by accident, especially in IIT programs that do not adopt a sterile insect technique approach, since fewer mosquitoes need to be released (Pagendam et al., 2020). An alternative way to increase the cost effectiveness of *Wolbachia* IIT programs is by expanding program coverage while keeping the quantity of mosquitoes constant through redistributing IIT mosquitoes from already-suppressed areas to new ones. While prior work has explored scale-down strategies in which reduced quantities of IIT mosquitoes were released, to our knowledge, no work has been done on either frequency-based scale-down strategies or redistribution strategies to expand program coverage, despite their practical importance.

Mathematical models have been used for studying mosquito population dynamics (Roques & Bonnefon, 2016) and the intervention efficacy of traditional vector control measures such as insecticide-based measures (Barbosa et al., 2018) and the sterile insect technique (Dufourd & Dumont, 2013). While mathematical models exist for studying *Wolbachia* IIT release optimization (Magori et al., 2009; Pagendam et al., 2020; Soh et al., 2022; Matsufuji & Seirin-Lee, 2023), these models cannot model large spatial scales and more importantly, do not consider exogenous factors affecting the intervention efficacies of *Wolbachia*-based programs such as precipitation, temperature, and mosquito migration. In addition to playing a significant role in the mosquito life cycle (Reinhold et al., 2018; Alto & Juliano, 2001), climatic factors such as temperature have been found to impact the intervention efficacy of *Wolbachia* replacement programs (Vásquez et al., 2023), and the same could hold for IIT programs. Prior evidence from randomized controlled trials and field trials also suggest that the migration of *Wolbachia* mosquitoes in treatment boundary areas has non-negligible effects on both IIT as well as replacement programs (Bansal et al., 2023; Utarini et al., 2021), which makes the characterization of mosquito population dynamics over both spatial and temporal scales important.

To address these issues, we propose a process-based model that explicitly accounts for precipitation, temperature, and spatio-temporal mosquito population dynamics. We employed the model to study the intervention efficacy of various release optimization strategies for an existing *Wolbachia* IIT program that targets *Aedes aegypti* mosquitoes. We used Singapore as the study site because it has an established network of mosquito surveillance traps which systematically collects national data on mosquito abundance (Ong et al., 2020), and extensive *Wolbachia* IIT field trial information for model validation purposes (The Project *Wolbachia* - Singapore Consortium & Ng, 2021; Bansal et al., 2023). Using the model, we sought to understand the consequences of:

i. Quantity- and frequency-reduction scale-down strategies in existing release areas to reduce the number of mosquitoes released, and
ii. Program expansion strategies that cover more release areas while keeping the overall mosquito release quantity constant, with the goal of comparing the trade offs between intervention efficacy and the program resources required to adopt each *Wolbachia* IIT release strategy, using Singapore as a case study.

## Methods

### Modeling mosquito metapopulation dynamics with a process-based framework

We developed a process-based framework, described as a system of ordinary differential equations that represents various life cycle stages of both *Wolbachia*-positive and *Wolbachia*-negative *Aedes aegypti* mosquitoes. This model was adapted from Cailly et al. (2012) and modified by adding new compartments to explicitly represent male mosquitoes, *Wolbachia*-positive mosquitoes, and mating-cross outcomes in adult female mosquitoes. Fluxes were also added to account for the migration of adult mosquitoes and introduction of *Wolbachia*-positive mosquitoes through the IIT program.

The 34-compartment model shown in Figure 1 represents *Wolbachia*-positive (*w*) and *Wolbachia*-negative (*u*) mosquitoes in the aquatic and adult life cycle compartments. The aquatic stages comprise the egg (*E*), larvae (*L*), pupae (*P*) compartments; for example, *P*_*u*_ represents the compartment for *Wolbachia*-negative pupae. The rates of change for the aquatic compartments are as follows:

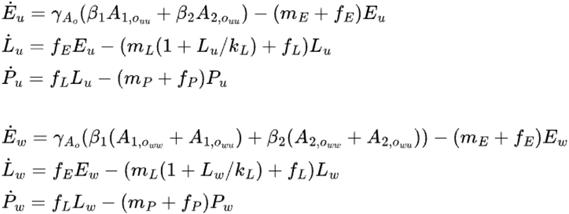

**Figure 1.**
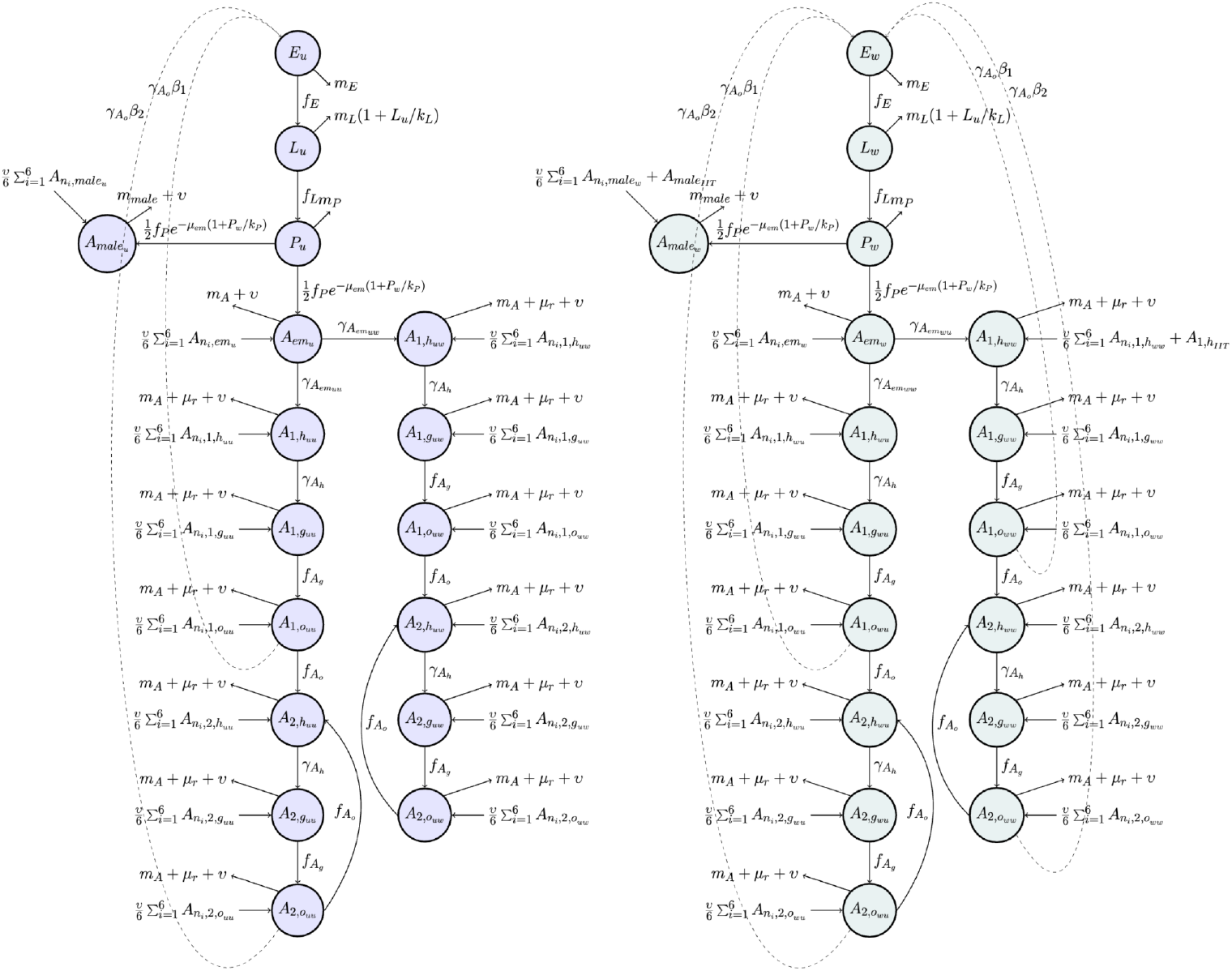
Model diagram for *Wolbachia*-negative (left) and *Wolbachia*-positive (right) mosquitoes across various life cycle stages.

**Figure 2.**
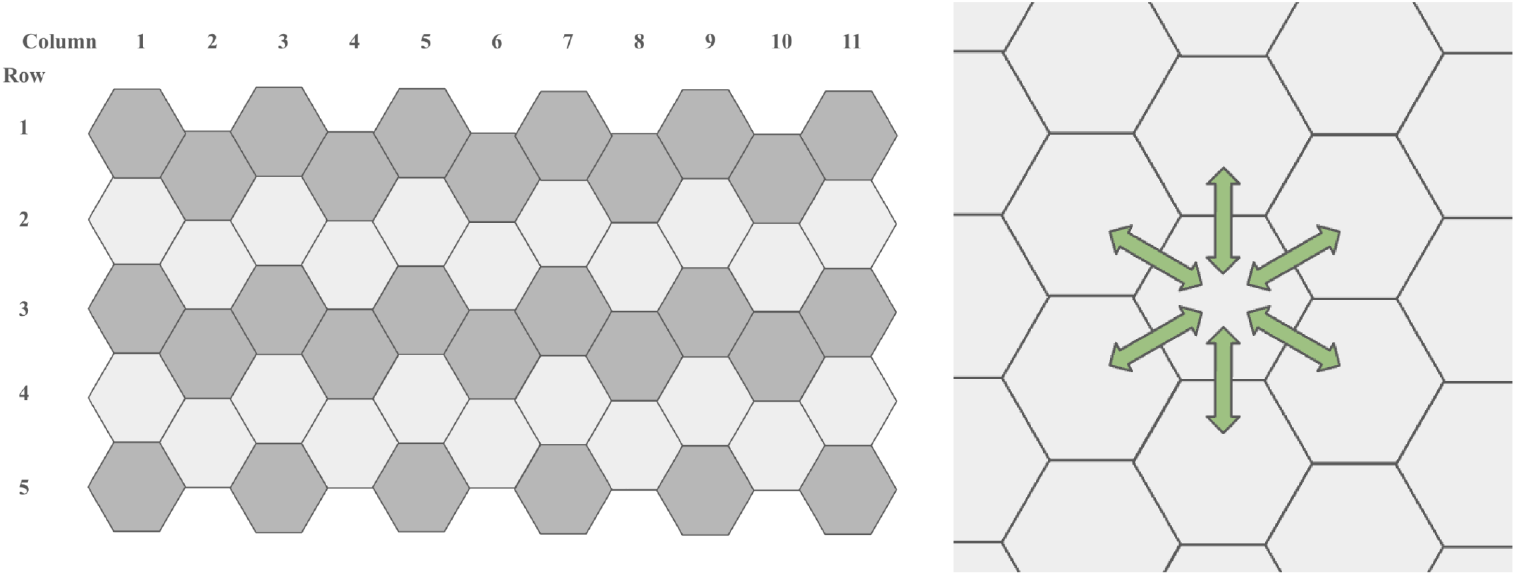
The spatial distribution of mosquitoes was represented by a grid of vertical hexagons. Mosquitoes mix freely within each hexagon (left). Some mosquitoes move between neighboring hexagons at each timestep (right).

**Figure 3.**
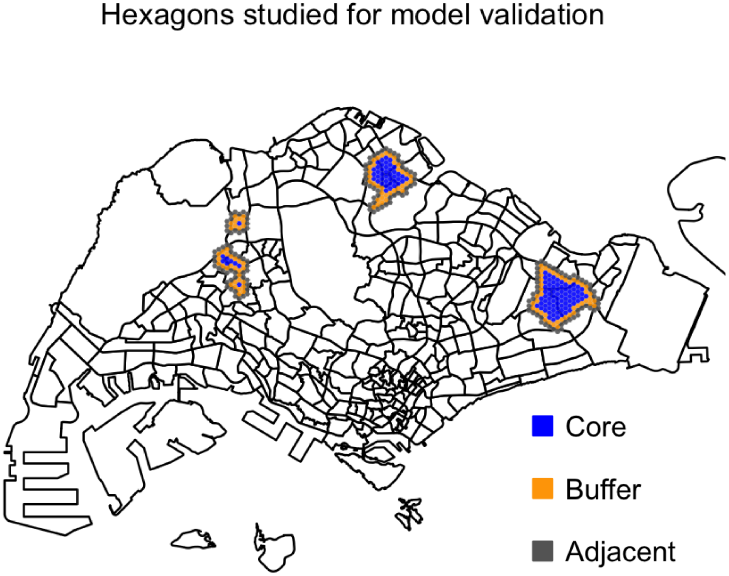
Release hexagons modeled after the early release program in Singapore. Core, buffer, and adjacent non-release hexagons are colored in blue, orange, and gray respectively.

**Figure 4.**
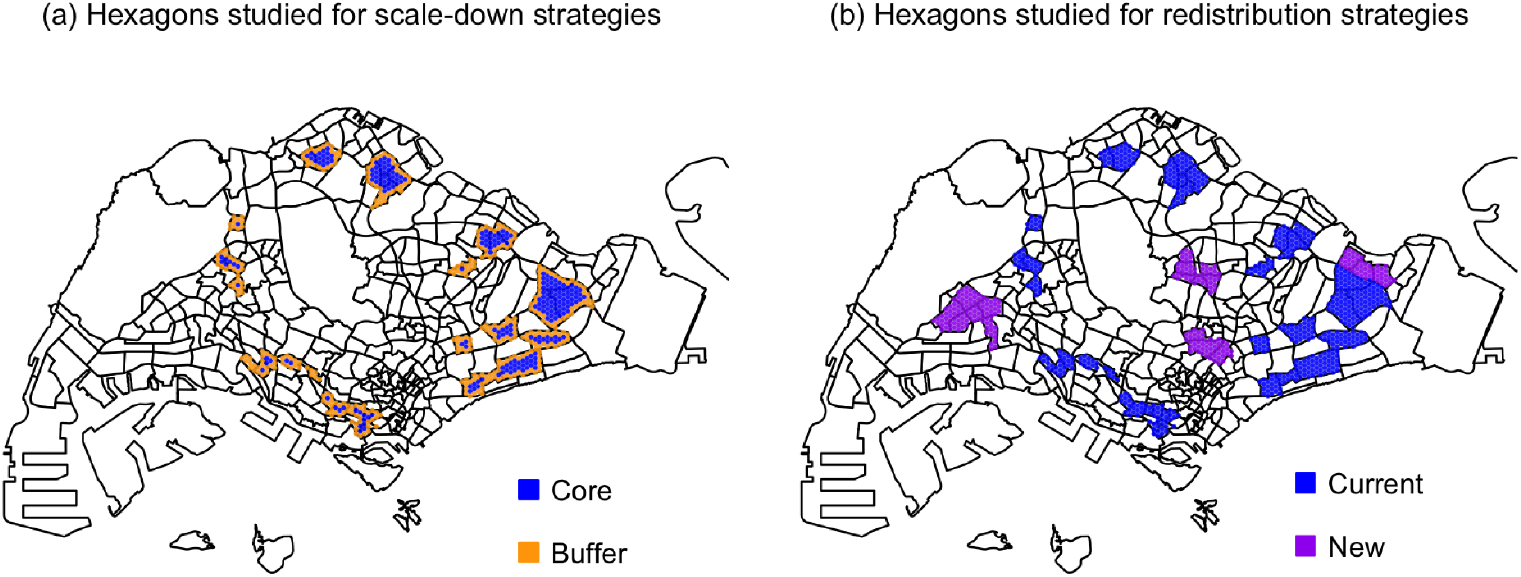
Release hexagons for the current release program in Singapore, core and buffer release hexagons colored in blue and orange respectively. (a). Release hexagons for the proposed expanded release program in Singapore, current and new release hexagons colored in blue and purple respectively (b).

**Figure 5.**
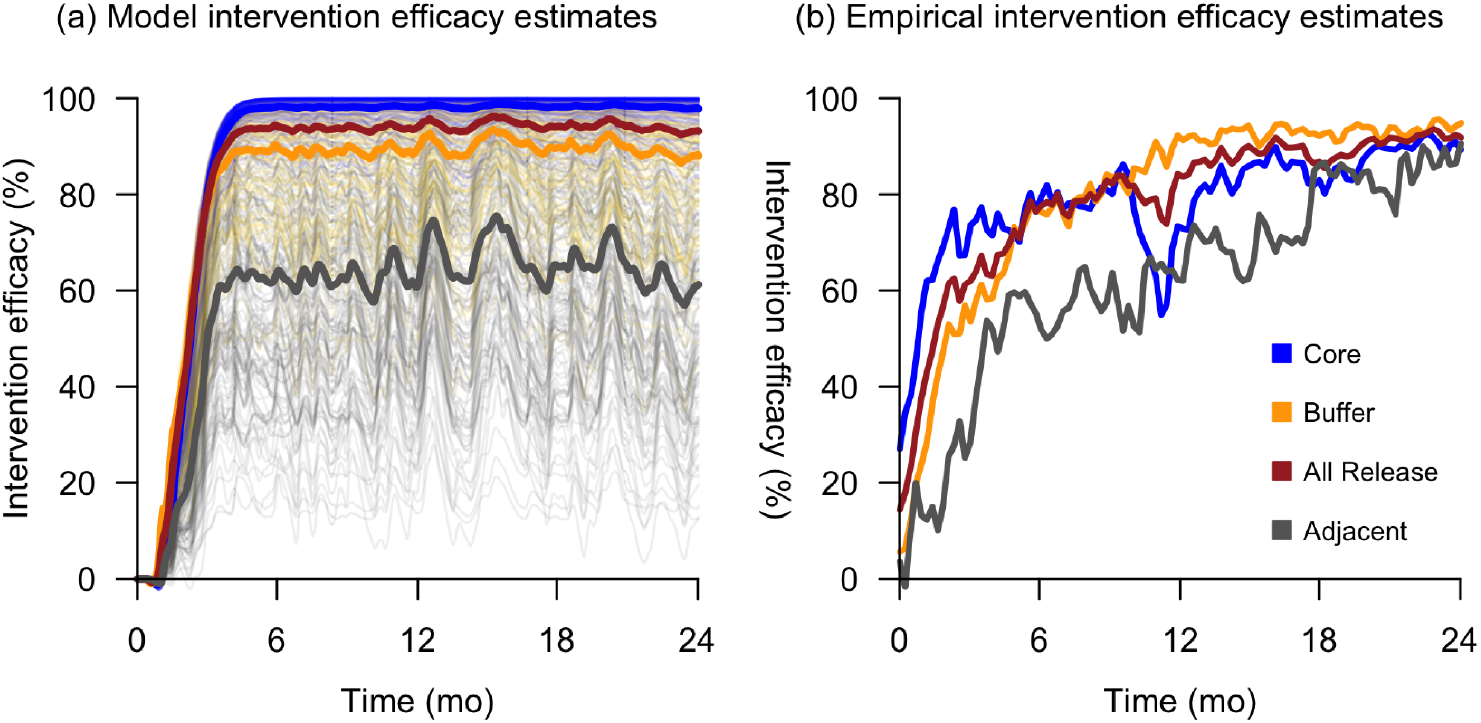
Model based simulation (a) and the empirically estimated (b) intervention efficacies of the early release program in Singapore. The intervention efficacy was measured by the difference in adult ovipositing female abundance between the intervention and baseline (no intervention). The x-axis represents time since the intervention began. Each thin line represents the intervention efficacy for a single hexagon, whereas each thick line represents the mean intervention efficacy for an entire hexagon class. Core, buffer, and adjacent non-release hexagons are colored in blue, orange, and gray respectively. The mean intervention efficacy line for all release hexagons (core and buffer) is shown in red.

**Figure 6.**
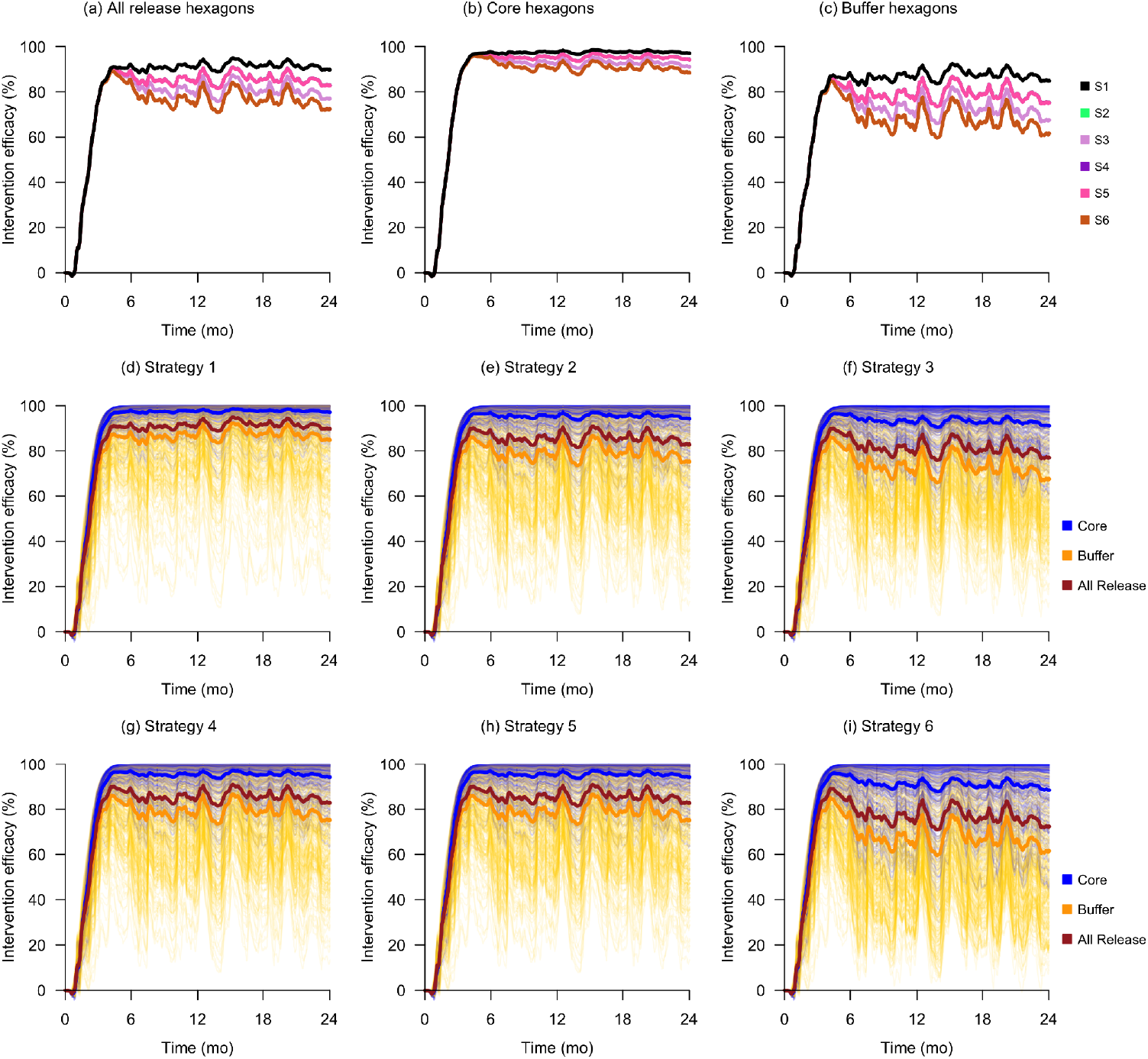
Intervention efficacy outcomes of various scale-down strategies. Plots (a), (b), and (c) show the mean intervention efficacy in all release, core-only, and buffer-only hexagons respectively for all strategies. The remaining plots (d) to (i) show the detailed intervention efficacies in all hexagons for Strategies 1 to 6 respectively.

**Figure 7.**
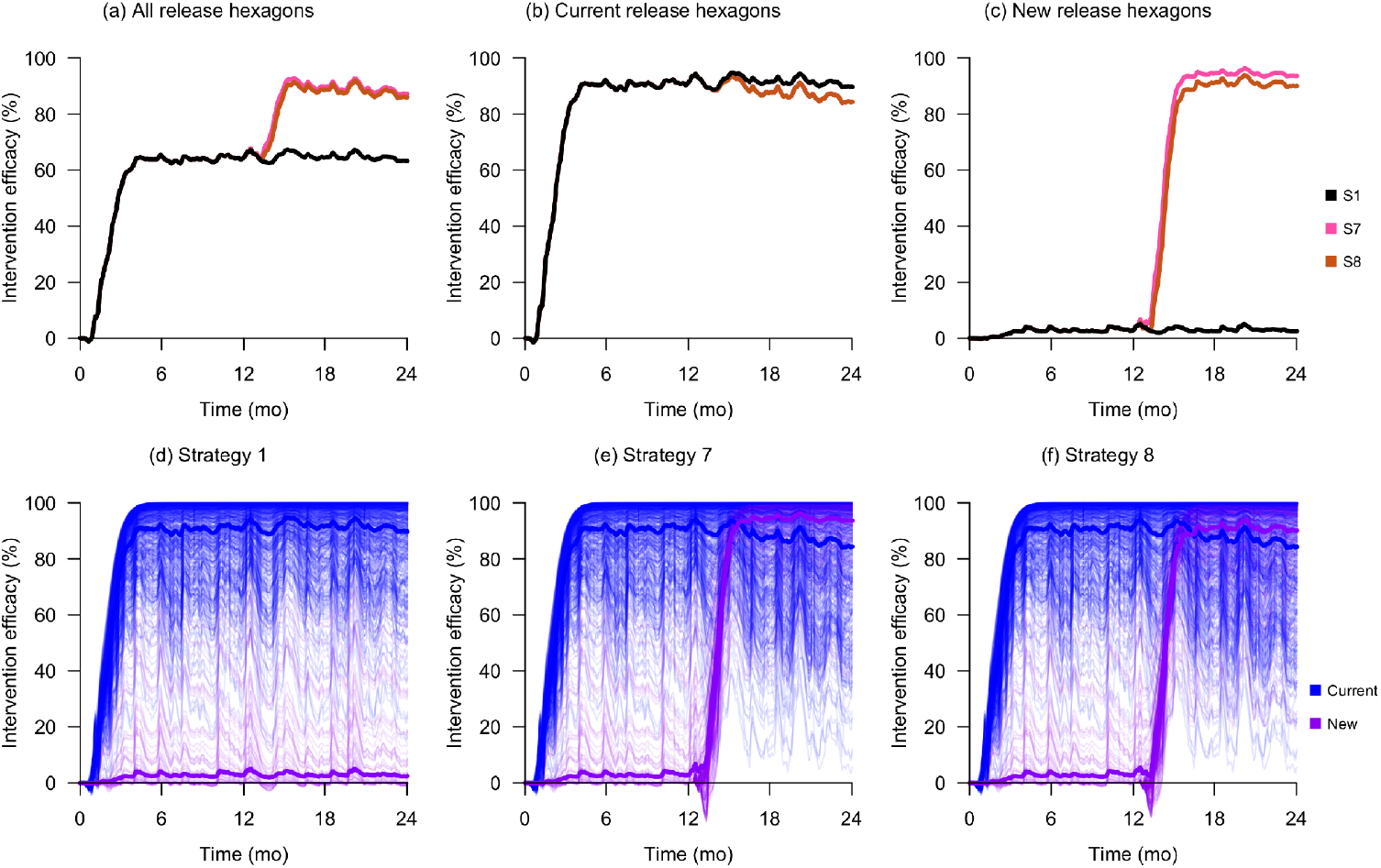
Intervention efficacy outcomes of various redistribution strategies. The first row of plots show the mean intervention efficacy in both current and new hexagons (a), current hexagons (b), and new hexagons only (c). The remaining plots (d), (e), and (f) show the intervention efficacies in all hexagons for Strategies 1, 7, and 8 respectively.

The adult stages are divided into emergent (*A*_*em*_), nulliparous (*A*_*1*_), and parous (*A*_*2*_) female adults, and male adults (*A*_*male*_). Nulliparous and parous female adults are further divided into host-seeking (*h*), resting (*g*), and ovipositing (*o*). Additionally, adult nulliparous and parous females are divided into their mating crosses (for example, *A*_*2,huw*_ represents nulliparous host-seeking *Wolbachia*-negative females that mated with a *Wolbachia*-positive male). The rates of change for the adult stages are represented as follows:

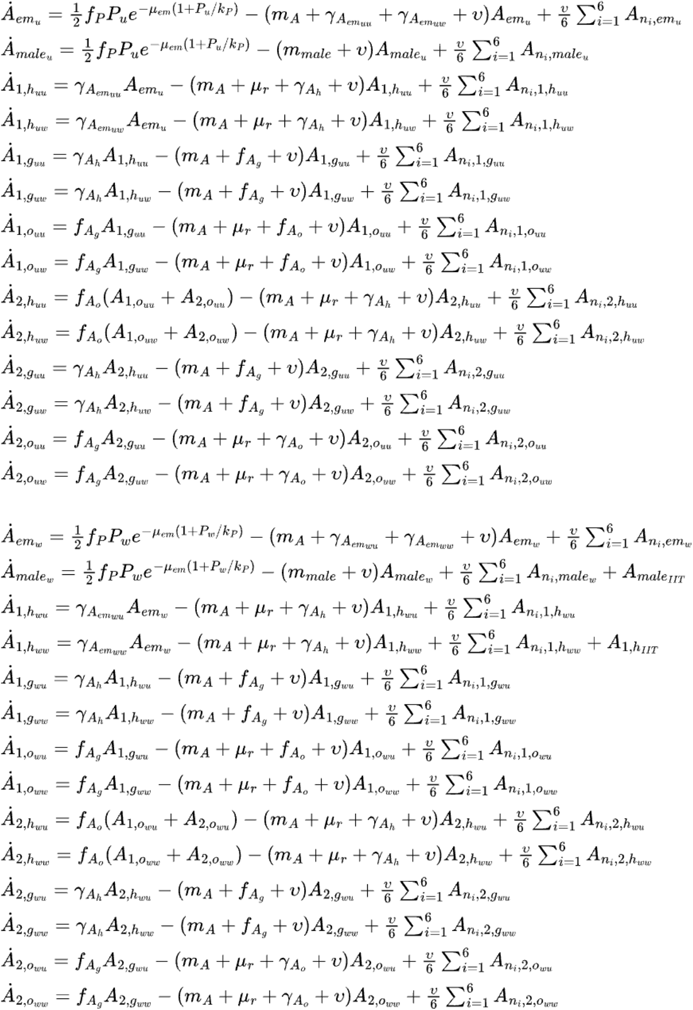

where

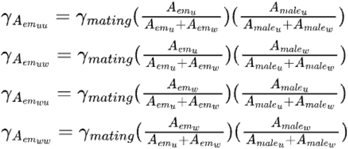

The parameter values and functions used in this model were either directly obtained from Bonnin et al. (2022) and Tran et al. (2020) or estimated from data in the literature. The parameter values and functions are described in detail in the Supplementary Material. For a given compartment *X, β*_*X*_ is the egg-laying rate, *γ*_*X*_ is the transition rate to the subsequent compartment, *μ*_*X*_ is the mortality rate, *μ*_*r*_ is the additional mortality rate associated with host-seeking behavior, and *υ* is the mosquito migration rate. The functions represent the time-varying temperature- and precipitation-dependent transition rates (*f*_*X*_), mortality rates (*m*_*X*_), and total carrying capacity indices (*k*_*X*_). The total carrying capacity is a sum of two components: a human-dependent (*k*_*Xfix*_) and a precipitation-dependent one (*k*_*Xvar*_). The human-dependent component represents the availability of man-made oviposition sites inside of homes such as stagnant water in flower pots and vases, whereas the precipitation-dependent component represents the availability of oviposition sites that occur due to rain.

Given the effects of adult mosquito migration on mosquito abundance and treatment program efficacy, the framework was designed to factor in migration by explicitly defining the study area as a grid of vertical hexagons. A single hexagon approximates a planning sector, which is the primary spatial resolution used for planning the surveillance and control of environment-related infectious diseases in Singapore (Bansal et al., 2023). Mosquitoes are modeled to mix homogeneously within a hexagon, and some adult mosquitoes migrate out of a hexagon into its six neighbors at a total daily rate of *υ*; conversely adult mosquitoes migrate into a hexagon from each of its six neighbors at a daily rate of *υ*/6. The number of mosquitoes in each of the neighboring hexagons is denoted by *A*_*n*_.

### Climate, geospatial, and human population data

In this study, we ran all simulations over a two-year period using the historical daily mean temperature and precipitation data obtained from the Meteorological Service Singapore website (*Historical Daily Records*, n.d.). Specifically, these data were obtained for the two-year period starting 1 January 2020 and ending 31 December 2021 from four weather stations across the country: Admiralty, Jurong West, Newton, and Paya Lebar. Each hexagon was assigned the set of climate data belonging to its nearest weather station. Additionally, a 300-day burn-in period was added prior to the start of each simulation using the mean values of the two-year climate dataset to allow mosquito abundances to reach equilibrium before interventions began.

The grid of vertical hexagons was designed to span the main island of Singapore, Pulau Ujong. Each hexagon was designed to have a side length of 184m to match the average planning sector area of 0.088km^2^ (Bansal et al., 2023). A hexagon was classified as a land or water one based on the geospatial subzone in which its center lies. The subzone boundary latitude and longitude coordinates were obtained from the Singapore government data portal (*Master Plan 2014 Subzone Boundary (No Sea) — Data*.*gov*.*sg*, n.d.). Land hexagons were assigned the maximum human- and precipitation-dependent carrying capacity indices of 100 and 50 respectively. Water hexagons were given 0 for both indices since mosquito breeding is taken not to occur there.

We assumed that the effective human-dependent mosquito carrying capacity was proportional to the number of residents in each hexagon. Subzone-level resident data from 2023 was obtained from the Singapore Department of Statistics (*Singapore Residents by Planning Area / Subzone, Age Group, Sex and Type of Dwelling, June 2023*, n.d.), and each hexagon was then assigned a normalized human-density value (*H*_*norm*_) by taking the number of residents in the subzone in which its center lies, and normalizing that against the number of residents in the most populous subzone.

### Model validation

Validation was performed by comparing model-based estimates of intervention efficacy against empirically-derived estimates in Bansal et al. (2023). Specifically, we ran a two-year simulation modeled after the early *Wolbachia* IIT release program conducted in four townships across Singapore from 2020 to 2022. We identified the approximate subzones belonging to the release townships based on publicly available release maps (*Wolbachia-Aedes Release Schedule*, n.d.), and used them to select release hexagons whose centers fell in the abovementioned subzones.

We studied the intervention efficacy in release hexagons as well as adjacent non-release hexagons. Release hexagons were classified into core or buffer, where buffer hexagons have at least one non-release neighboring hexagon, while core hexagons do not have any non-release neighboring hexagons. Adjacent non-release hexagons were defined to have at least one release neighboring hexagon. The intervention efficacy was defined as the percentage reduction of adult ovipositing female mosquito abundance between the intervention and the baseline where no intervention occurred. Wild-type adult ovipositing female mosquito abundance was used as the empirical study used data from Gravitraps, which trap gravid females in search of oviposition sites (Ong et al., 2020).

Two simulations were run: a baseline simulation where no releases occurred, while the intervention simulation had releases twice a week at an initial overflooding ratio of 10:1. The initial overflooding ratio is defined as the ratio of *Wolbachia* IIT male mosquitoes released at each release event relative to the adult wild-type males at the start of the intervention. An initial overflooding ratio of 10:1 was chosen to mirror the mean of the two overflooding ratios (5:1 and 15:1) studied by Pagendam et al. (2020).

### Scale-down and redistribution strategies

We studied two classes of release strategies: scale-down strategies that optimize the current release program, and redistribution strategies that expand geographical program coverage without requiring more *Wolbachia* IIT mosquitoes. Each strategy is detailed in Table 3.

**Table 1.** Definition of scale-down and redistribution strategies.

| Strategy | Type | Name | Initial intervention condition | Threshold(s) to switch to new | New intervention condition(s) |
| --- | --- | --- | --- | --- | --- |
| 0 | - | Baseline | No intervention | - | - |
| 1 | - | Constant | Release in current treatment areas at the initial overflooding ratio of 10:1 | - | - |
| 2 | Scale-down | Half quantity | Release in current treatment areas at the initial overflooding ratio of 10:1 | When wild-type adult male abundance reaches 50% of initial | Release in current treatment areas at an initial overflooding ratio of 5:1 twice a week |
| 3 | Scale-down | Quarter quantity | Release in current treatment areas at the initial overflooding ratio of 10:1 | When wild-type adult male abundance reaches 50% and 25% | Release in current treatment areas at initial overflooding ratios of 5:1 and 2.5:1 twice a week respectively |
| 4 | Scale-down | Weekly | Release in current treatment areas at the initial overflooding ratio of 10:1 | When wild-type adult male abundance reaches 50% of initial | Release in current treatment areas at the initial overflooding ratio of 10:1 once a week |
| 5 | Scale-down | Alternate weeks | Release in current treatment areas at the initial overflooding ratio of 10:1 | When wild-type adult male abundance reaches 50% of initial | Release in current treatment areas at the initial overflooding ratio of 10:1 twice every alternate week |
| 6 | Scale-down | Fortnightly | Release in current treatment areas at the initial overflooding ratio of 10:1 | When wild-type adult male abundance reaches 50% of initial | Release in current treatment areas at the initial overflooding ratio of 10:1 once every two weeks |
| 7 | Redistribution | Redistribution by quantity | Release in current treatment areas at an initial overflooding ratio of 10:1 twice a week for the first year | After one year | Release in:<br>- Current treatment areas at an initial overflooding ratio of 5:1 twice a week for the second year.<br>- New treatment areas at an initial overflooding ratio of 10:1 |
| 8 | Redistribution | Redistribution by frequency | Release in current treatment areas at an initial overflooding ratio of 10:1 twice a week for the first year | After one year | Release in:<br>- Current treatment areas at an initial overflooding ratio of 10:1 once a week for the second year.<br>- New treatment areas at an initial overflooding ratio of 10:1 |

**Table 2.** Comparison of intervention efficacy and resource consumption for various scale-down strategies.

| Strategy | Name | Average mean intervention efficacy over the final 24 weeks (%), (95% CI) |  |  | Relative resource consumption change (%) |  |
| --- | --- | --- | --- | --- | --- | --- |
|  |  | Core | Buffer | All release | Number of *Wolbachia* mosquitoes released | Number of release events |
| 1 | Constant | 98 (87–100) | 87 (55–100) | 91 (60–100) | Reference | Reference |
| 2 | Half quantity | 95 (78–100) | 79 (42–99) | 85 (46–100) | -45 | 0 |
| 3 | Quarter quantity | 93 (71–100) | 72 (39–98) | 80 (41–100) | -65 | 0 |
| 4 | Weekly | 95 (78–100) | 79 (42–99) | 85 (46–100) | -45 | -45 |
| 5 | Alternate weeks | 95 (78–100) | 79 (42–99) | 85 (46–100) | -45 | -45 |
| 6 | Fortnightly | 90 (60–100) | 66 (28–98) | 76 (30–100) | -67 | -67 |

CI: Confidence interval

**Table 3.** Comparison of intervention efficacy and resource consumption for various redistribution strategies.

| Strategy | Name | Average mean intervention efficacy over the final 24 weeks (%), (95% CI) |  |  | Relative resource consumption (% change) |  |
| --- | --- | --- | --- | --- | --- | --- |
|  |  | Current | New | All release | Number of *Wolbachia* mosquitoes released | Number of release events |
| 1 | Constant | 91 (60–100) | 3 (0–42) | 65 (0–100) | Reference | Reference |
| 7 | Redistribution by quantity | 87 (46–100) | 94 (64–100) | 89 (47–100) | -7 | 22 |
| 8 | Redistribution by frequency | 87 (46–100) | 91 (48–100) | 88 (46–100) | -16 | -14 |

CI: Confidence interval

The *Wolbachia* IIT release program in Singapore was expanded to cover fifteen townships in 2024. We simulated five scale-down release strategies to optimize the current release program, which all start with an initial overflooding ratio of 10:1. To compare intervention efficacies for each release optimization strategy, release maps were used to identify release hexagons of these townships using the same method as per model validation. Strategy 1 is a constant release strategy modeled after the current program where IIT male mosquitoes are released twice a week. Strategies 2 and 3 are quantity reduction strategies where the number of released IIT male mosquitoes is reduced when the wild-type male abundance falls below specific thresholds. Strategies 4 to 6 are frequency reduction strategies where the number of release events is reduced when the wild-type male abundance falls below specific thresholds.

Given the possibility of reducing the number of mosquitoes released in already-suppressed hexagons without significantly affecting intervention efficacy, we also explored two redistribution strategies to increase program coverage without changing the overall number of mosquitoes produced. This was done by taking some mosquitoes from current release hexagons and releasing them in new hexagons belonging to four large townships. New hexagons were selected based on subzones with substantial population sizes, ensuring that the combined population of the newly released subzones was approximately half of that already covered by the current release program. Both Strategies 7 and 8 adopted the same constant release strategy for the first year in current release hexagons, and new hexagons did not receive any mosquitoes in the first year.

Strategy 7 is a redistribution-by-quantity strategy where in the second year, the quantity of mosquitoes released in the current release hexagons is reduced by half, but the release frequency is kept constant at twice a week; new hexagons receive redistributed mosquitoes twice a week at the initial overflooding ratio of 10:1. Strategy 8 is a redistribution-by-frequency strategy where in the second year, the frequency of mosquito releases in the current release hexagons is reduced to once a week, while the release quantity is kept constant at the initial value; new hexagons receive redistributed mosquitoes once a week at the initial overflooding ratio of 10:1.

The intervention efficacy was computed as the percentage reduction in wild-type adult female mosquito abundance for each strategy compared to the baseline simulation where no releases were conducted. We analyzed the average mean intervention efficacies over the final 24 weeks of the simulation for all release hexagons (*IE*_*final*_) and the resource consumption footprint for each strategy, specifically the total number of mosquitoes released and the total number of release events required for the entire two-year program.

## Results

### Model validation

The intervention efficacy estimated by the model outputs was compared against the empirically-derived estimates in Bansal et al. (2023). The model captured the dynamics observed in the early stages of the field trial: a larger suppression effect was observed in core hexagons than buffer hexagons, and non-negligible suppression effects were observed in adjacent non-release hexagons. Additionally, the model outputs and field trial estimates both reached >90% mean intervention efficacy at the end of the two-year intervention period. However, the model outputs differed from the trial estimates as time went on. Specifically, the field trial intervention efficacy in the core release areas fell below that of the buffer release areas in the latter half, and intervention efficacies across all release and adjacent non-release areas converged towards the end of the two-year period. We discuss the possible reasons for these deviations in detail later in the discussion section.

### Scale-down strategies to optimize the current release program

As expected, the constant release strategy (Strategy 1) had the best suppression with *IE*_*final*_ values of 91 % across all release hexagons. The half quantity (Strategy 2) and quarter quantity (Strategy 3) release strategies had *IE*_*final*_ values of 85% and 80% in all release hexagons but required approximately 45% and 65% fewer IIT mosquitoes respectively. The weekly (Strategy 4) and alternate weeks (Strategy 5) strategies both had *IE*_*final*_ values of 85% in all release hexagons and required 45% fewer IIT mosquitoes and release events. The fortnightly strategy (Strategy 6), on the other hand, produced a markedly lower *IE*_*final*_ value of 76% across all release hexagons, but required approximately 67% fewer IIT mosquitoes and release events.

### Redistribution strategies to increase program coverage

We analyzed the *IE*_*final*_ values and resource consumption of each program expansion strategy in the same manner as in the scale-down strategies, described in the previous section. Both the redistribution by quantity (Strategy 7) and redistribution by frequency (Strategy 8) strategies had better overall mean intervention efficacies across current and new hexagons than the constant release strategy (Strategy 1) because they led to minimal decrease in suppression for current release hexagons while creating a strong suppression effect in the new hexagons. Strategies 7 and 8 had overall *IE*_*final*_ values of 89% and 88% respectively, compared to 65% in Strategy 1. In current hexagons, Strategy 1 produced an *IE*_*final*_ of 91%, which was only very slightly higher than 87% in both Strategies 7 and 8. However, in new hexagons, Strategy 1 produced an *IE*_*final*_ of only 3%, whereas Strategies 7 and 8 resulted in much higher overall *IE*_*final*_ values of 94%, and 91% respectively. In terms of resource consumption, although Strategy 7 required 7% fewer mosquitoes than Strategy 1, it required 22% more release events; Strategy 8 on the other hand, required approximately 16% fewer mosquitoes and 14% fewer release events.

## Discussion

As the *Wolbachia* incompatible insect technique becomes a more commonly-used tool to combat mosquito-borne diseases across various regions in the world, there is a growing need to optimize release programs to maximize cost-effectiveness. Given the potential risks of releasing too few IIT mosquitoes, it is pertinent to use mathematical models to explore optimal release strategies beforehand. While there exist mathematical models to study release strategies, they are unable to explore IIT programs over large spatial scales. Models by Pagendam et al. (2020), Soh et al. (2022), and Matsufuji & Seirin-Lee (2023) do not take into account important climatic factors affecting the life cycle of mosquitoes such as precipitation and temperature. Additionally, the models by Soh et al. (2022) and Matsufuji & Seirin-Lee (2023) assume a well-mixed ecological environment, which does not account for heterogeneous geographical release coverage in real *Wolbachia* IIT programs. While an agent-based model proposed by Magori et al. (2009) accounts for climatic factors, it requires extremely fine-grained descriptions of individual houses and breeding containers as well as tracks individual mosquitoes, making it extremely challenging for studying IIT programs over a large city. Therefore, it was important for us to develop a general model that could explore program resource requirements over large spatial scales while accounting for precipitation, temperature and mosquito migration.

In this study, simulations of various scale-down and redistribution strategies were performed using a process-based model to optimize an existing *Wolbachia-Aedes aegypti* IIT program in Singapore. To our knowledge, no studies have been done so far to explore redistribution strategies or frequency-based scale-down strategies. Pagendam et al. (2020) explored two quantity-based scale-down strategies using a stochastic model and determined that at low female contamination rates, one of them (the crude adaptive strategy) was able to completely eliminate wild-type mosquitoes at a very high probability, almost identical to that of the constant release strategy. We explored a similar strategy with our model (the quarter quantity release strategy, Strategy 3) and found that while there was still effective suppression in release areas, the mean intervention efficacy across all release hexagons fell noticeably by 11% (from 91% to 80%) compared to the constant release strategy (Strategy 1). We posit that the appreciable decrease observed in the simulation outputs was because our model factored in migratory effects of mosquitoes at the buffer hexagons; we observed that the *IE*_*final*_ for core hexagons only fell by 5% (from 98% to 93%), whereas the decrease was 15% (from 87% to 72%) for buffer hexagons. These results highlight the dilutive effects of mosquito migration on intervention efficacy at release boundary areas, which is corroborated by field trial results in both *Wolbachia* IIT and replacement programs (Bansal et al., 2023; Utarini et al., 2021).

In general, our simulations identified a few quantity- and frequency-based scale-down strategies that could lead to significantly less resource consumption while having minimal impact on the overall mean intervention efficacy of *Wolbachia* IIT programs. For example, the half quantity (Strategy 2) and weekly (Strategy 4) release strategies resulted in a small decrease in overall intervention efficacy of 6% (from 91% to 85%) compared to the constant release strategy (Strategy 1). This is despite them requiring significantly fewer resources compared to Strategy 1; Strategies 2 and 4 both required 45% fewer mosquitoes, and Strategy 4 additionally reduced the number of release events by 45%. Given that they have similar overall intervention outcomes, reducing the release frequency from twice to only once a week would lead to the most cost-effective program outcomes especially if release events contribute significantly to program costs.

Simulations of redistribution strategies also showed that both strategies led to significantly higher overall intervention efficacy compared to the constant release strategy (Strategy 1), because they caused strong suppression in new release areas while having minimal reduction in intervention efficacy in existing release areas. The redistribute-by-quantity (Strategy 7) strategy, however, required a 22% increase in release events compared to the constant release strategy (Strategy 1), whilst the redistribute-by-frequency (Strategy 8) strategy required 16% fewer mosquitoes and 14% fewer release events. Since both redistribution strategies have similar intervention outcomes, the redistribute-by-frequency strategy is the most cost-effective, since it would result in fewer mosquitoes and release events whilst having greater intervention efficacy than the existing program.

While the simulation results suggest that strategies to reduce the release frequency in both scale-down and redistribution strategies will significantly reduce program resources while maintaining strong suppression, it is also important to consider them in the context of other existing mosquito control measures that could affect the released *Wolbachia* mosquitoes. For example, the actual intervention efficacy of the strategy could be affected by the frequency and timing of existing insecticide-spraying activities in release areas.

Therefore, the recommended strategies should be implemented in a phased approach. First, a few release areas with strong suppression should be selected to adopt a scale-down strategy and monitored over a period of time. If suppression remains effective over an extended period of time, the excess mosquitoes that were produced for these areas can then be released in new areas using one of the redistribution strategies. Then, both existing and new release areas that have adopted the new release strategy should then be observed for a period of time. If there remains effective suppression in both existing and new areas, these steps can subsequently be repeated for the other existing release areas.

### Limitations

There were deviations between the model outputs and the empirical intervention efficacy estimates, possibly due to complexities that arise in the conduct of an actual *Wolbachia* release program. The model simulations were run with the assumption that the number of mosquitoes released remained constant throughout the two-year period and the core and buffer hexagons remained the same due to the lack of data availability. However, in reality, the release program began with a smaller group of sites and expanded over time (Bansal et al., 2023). This meant that release sites switched identities from adjacent non-release to buffer, and buffer to core sites over time in the empirical study. Therefore, the aggregation of intervention efficacy by event time was also confounded by anthropogenic and environmental characteristics across different locations and time points due to the staggered introduction of *Wolbachia* releases in real sites. Consequently, we observed some suppression in the empirical intervention efficacy estimates even at the start of the field trial period across release and adjacent non-release areas. It should be noted that the suppression effect observed at the start of the trial period could also be due to the fact that although the field trial analyzed data from 2020 to 2022, *Wolbachia* releases had already begun in Singapore in 2016 (Bansal et al., 2023). Moreover, the empirical results were compared against those from counterfactual locations in the same time period using the synthetic control method, whereas our study used outputs from the simulation where no releases had occurred as the baseline for comparison.

### Further applications of the spatiotemporal mosquito metapopulation model

As the model was designed to be flexible and configurable with various climate and geospatial inputs, we foresee its application in further *Wolbachia* IIT-based studies. For example, the model can be used in the planning and optimization of new and existing *Wolbachia* IIT release programs in a variety of regions around the world, including those with subtropical and temperate climates with seasonal *Aedes aegypti* population fluctuations (Mores et al., 2020; Benitez et al., 2021). Additionally, the existing model can also be configured to study and optimize *Wolbachia* replacement programs where both adult male and female *Wolbachia* mosquitoes are released, or modified to model replacement programs where mosquitoes are released at different lifecycle points such as the egg stage (Allman et al., 2023).

Modeling studies have been conducted on the effects of climate change on *Wolbachia* replacement programs (Vásquez et al., 2023). Similarly, our model can be used to predict the effects of climate change on existing *Wolbachia* IIT release programs, and study various strategies to make such programs more resilient to future climatic variations. The model can also easily be configured to model the population dynamics and *Wolbachia* IIT-based control of other mosquito species such as *Aedes albopictus*, which is becoming an increasingly important vector in Europe because of their potentially expanding range due to climate change (Oliveira et al., 2021).

Finally, given that the keen interest in mosquito population dynamics stems from its implications for mosquito-borne disease burden, we expect that our model can also be used to study the intervention efficacy and cost effectiveness of various *Wolbachia* IIT program strategies on disease dynamics and burden. Studies have been done in the past where outputs from the mosquito dynamics model were used as inputs to mosquito-borne disease transmission models (Matsufuji & Seirin-Lee, 2023). Our model therefore provides the ability to tie *Wolbachia* IIT program inputs to mosquito population dynamics and eventually human disease burden.

## Conclusions

We developed a new modeling tool to study *Wolbachia* IIT programs over large spatial scales that explicitly accounts for climatic factors such as precipitation and rainfall as well as the movement of mosquitoes. Several scale-down and redistribution strategies were simulated to understand the trade-offs between intervention efficacy and program resource consumption. Results from our study suggest that both scale-down and redistribution strategies can be adopted to significantly reduce the resources required by *Wolbachia* IIT programs without compromising on intervention efficacy.

## Acknowledgements

The opinions expressed in this article are those of the individual authors, and do not reflect those of the primary author’s employer (Verily Life Sciences LLC). Verily Life Sciences produces *Wolbachia* mosquitoes to reduce mosquito populations.

