## Supplementary Material for "*Wolbachia* incompatible insect technique program optimization over large spatial scales using a process-based model of mosquito metapopulation dynamics"

**Table S1.** Process-based model parameters.

| Notation | Definition | Value | Reference |
|---|---|---|---|
| $\beta_1$ | Number of eggs laid per ovipositing nulliparous female | 60.0 | Bonnin et al., 2022 |
| $\beta_2$ | Number of eggs laid per ovipositing parous female | 80.0 | Bonnin et al., 2022 |
| $\gamma_{mating}$ | Transition rate from emergent to host-seeking adults per day | 0.4 | Bonnin et al., 2022 |
| $\gamma_{Ah}$ | Transition rate from host-seeking to engorged adults per day | 0.2 | Bonnin et al., 2022 |
| $\gamma_{Ao}$ | Minimum transition rate from ovipositing to host-seeking adults per day | 0.2 | Bonnin et al., 2022 |
| $\mu_{\mathrm{E}}$ | Minimum egg mortality rate per day | 0.01 | Bonnin et al., 2022 |
| $\mu_{\mathrm{em}}$ | Mortality rate during emergence per day | 0.1 | Bonnin et al., 2022 |
| $\mu_{\mathrm{r}}$ | Mortality rate related to seeking behavior per day | 0.08 | Bonnin et al., 2022 |
| $k_{Lfix}$ | Maximum human-dependent larval carrying capacity index in a specific hexagon | 100 | Reference index |
| $k_{Pfix}$ | Maximum human-dependent larval carrying capacity index in a specific hexagon | 100 | Reference index |
| $k_{Lvar}$ | Maximum rainfall-dependent larval carrying capacity index in a specific hexagon | 50 | Estimated |
| $k_{Pvar}$ | Maximum rainfall-dependent pupal carrying capacity index in a specific hexagon | 50 | Estimated |
| $\upsilon$ | Migration rate of adult mosquitoes out of a hexagon per day | 0.0663 | Estimated |

**Table S2.** Process-based model functions.

| Notation | Definition | Expression | Reference |
|---|---|---|---|
| $f_E$ | Transition function from egg to larva | $$\frac{24\rho\,\frac{T^K}{298}e^{\frac{\Delta H_A}{R}(\frac{1}{298}-\frac{1}{T^K})}}{1+e^{\frac{\Delta H_H}{R}(\frac{1}{T_{1/2H}}-\frac{1}{T^K})}}$$ Where $\rho = 0.01066$; $\Delta H_A = 10{,}798.18$; $\Delta H_H = 100{,}000$; $T_{1/2H} = 14{,}184.5$ | Bonnin et al., 2022 |
| $f_L$ | Transition function from larva to pupa | $$\frac{24\rho\,\frac{T^K}{298}e^{\frac{\Delta H_A}{R}(\frac{1}{298}-\frac{1}{T^K})}}{1+e^{\frac{\Delta H_H}{R}(\frac{1}{T_{1/2H}}-\frac{1}{T^K})}}$$ Where $\rho = 0.00873$; $\Delta H_A = 26{,}018.51$; $\Delta H_H = 55{,}990$; $T_{1/2H} = 304.58$ | Bonnin et al., 2022 |
| $f_P$ | Transition function from pupa to emergent adult | $$\frac{24\rho\,\frac{T^K}{298}e^{\frac{\Delta H_A}{R}(\frac{1}{298}-\frac{1}{T^K})}}{1+e^{\frac{\Delta H_H}{R}(\frac{1}{T_{1/2H}}-\frac{1}{T^K})}}$$ Where $\rho = 0.0161$; $\Delta H_A = 14{,}931.94$; $\Delta H_H = -472{,}379$; $T_{1/2H} = 148.45$ | Bonnin et al., 2022 |
| $f_{Ag}$ | Transition function from engorged adult to ovipositing site-seeking female adult | $$\frac{24\rho\,\frac{T^K}{298}e^{\frac{\Delta H_A}{R}(\frac{1}{298}-\frac{1}{T^K})}}{1+e^{\frac{\Delta H_H}{R}(\frac{1}{T_{1/2H}}-\frac{1}{T^K})}}$$ Where $\rho = 0.00898$; $\Delta H_A = 15{,}725.23$; $\Delta H_H = 1{,}756{,}481.07$; $T_{1/2H} = 447.17$ | Bonnin et al., 2022 |
| $f_{Ao}$ | Transition function from ovipositing adult to host-seeking female adult | $$\gamma_{Ao} \times (1 + P_{norm})$$ | Bonnin et al., 2022 |
| $m_E$ | Egg mortality rate | $\mu_E + 0.1$ if $P > 80$, $\mu_E$ otherwise | Bonnin et al., 2022 |
| $m_L$ | Larval mortality rate | $0.52 + 0.0007e^{0.1838(T-10)}$ if $P > 80$, $0.02 + 0.0007e^{0.1838(T-10)}$ otherwise | Bonnin et al., 2022 |
| $m_P$ | Pupal mortality rate | $0.52 + 0.0003e^{0.2228(T-10)}$ if $P > 80$, $0.02 + 0.0003e^{0.2228(T-10)}$ otherwise | Bonnin et al., 2022 |

| | | | |
|---|---|---|---|
| $m_A$ | Adult female mortality rate | $0.025 + 0.0003e^{0.1745(T-10)}$ | Bonnin et al., 2022 |
| $m_{male}$ | Adult male mortality rate | $0.05 + 0.0006e^{0.1745(T-10)}$ | Estimated |
| $k_L$ | Total larval carrying capacity index per hexagon | $k_L = H_{norm} \times k_{Lfix} + P_{norm} \times k_{Lvar}$ | Adapted from Tran et al., 2020 |
| $k_P$ | Total pupal carrying capacity index per hexagon | $k_P = H_{norm} \times k_{Pfix} + P_{norm} \times k_{Pvar}$ | Adapted from Tran et al., 2020 |

Note: $H_{norm}$ is the number of people living in the subzone in which the center of the hexagon falls, normalized against the most populated subzone to vary from 0 to 1. $P_{norm}$ is the amount of cumulative rainfall over one week and normalized to vary from 0 to 1.

**Sensitivity Analyses**

We performed sensitivity analyses of the mean $IE_{final}$ values for each strategy by varying the key model parameters by 30% above and below the main experiment value. Three parameter values (male mortality, overflooding ratio, and migration rate) were given a wider range of values.

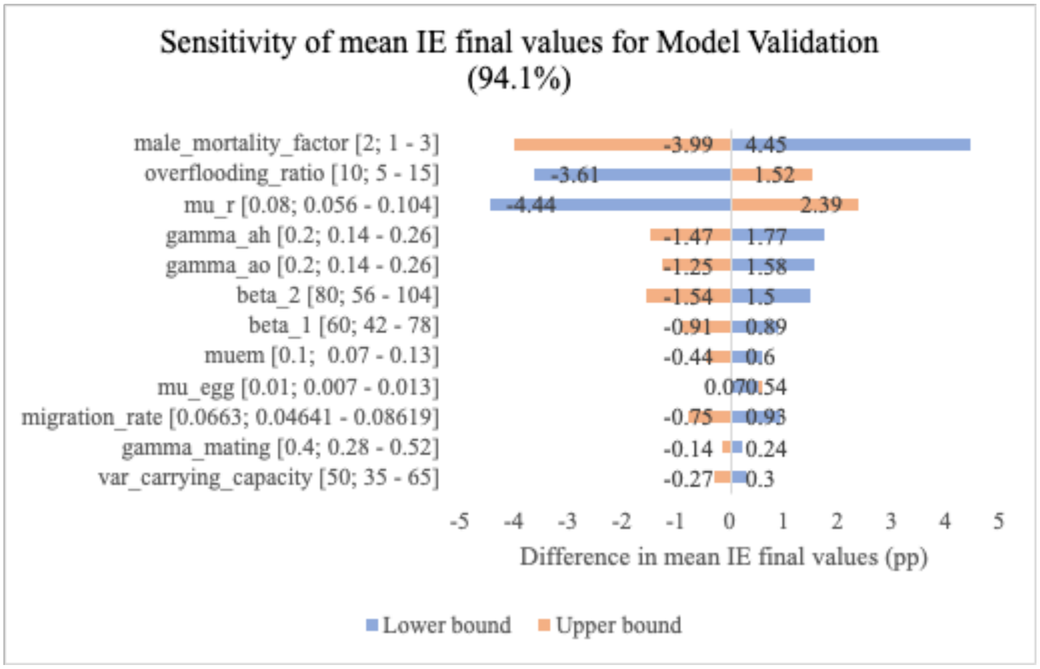

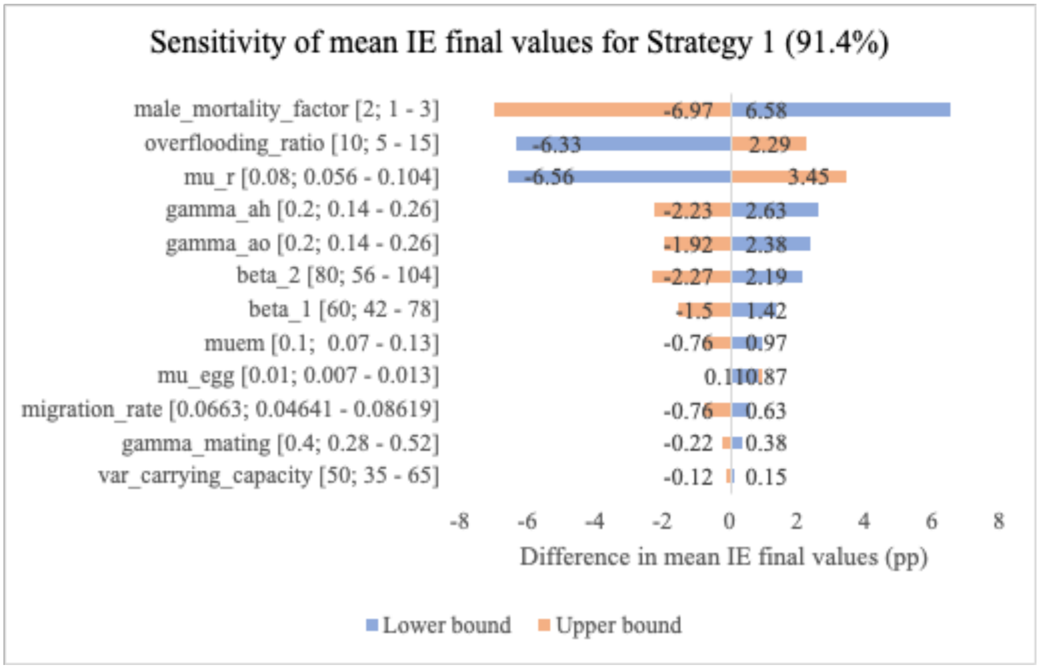
Sensitivity of mean IE final values for Strategy 1 (91.4%)
male_mortality_factor [2; 1 - 3]
overflooding_ratio [10; 5 - 15]
mu_r [0.08; 0.056 - 0.104]
gamma_ah [0.2; 0.14 - 0.26]
gamma_ao [0.2; 0.14 - 0.26]
beta_2 [80; 56 - 104]
beta_1 [60; 42 - 78]
muem [0.1; 0.07 - 0.13]
mu_egg [0.01; 0.007 - 0.013]
migration_rate [0.0663; 0.04641 - 0.08619]
gamma_mating [0.4; 0.28 - 0.52]
var_carrying_capacity [50; 35 - 65]
-6.97
6.58
-6.33
2.29
-6.56
3.45
-2.23
2.63
-1.92
2.38
-2.27
2.19
-1.5
1.42
-0.76
0.97
-0.76
0.63
-0.22
0.38
-0.12
0.15
-8
-6
-4
-2
0
2
4
6
8
Difference in mean IE final values (pp)
Lower bound
Upper bound

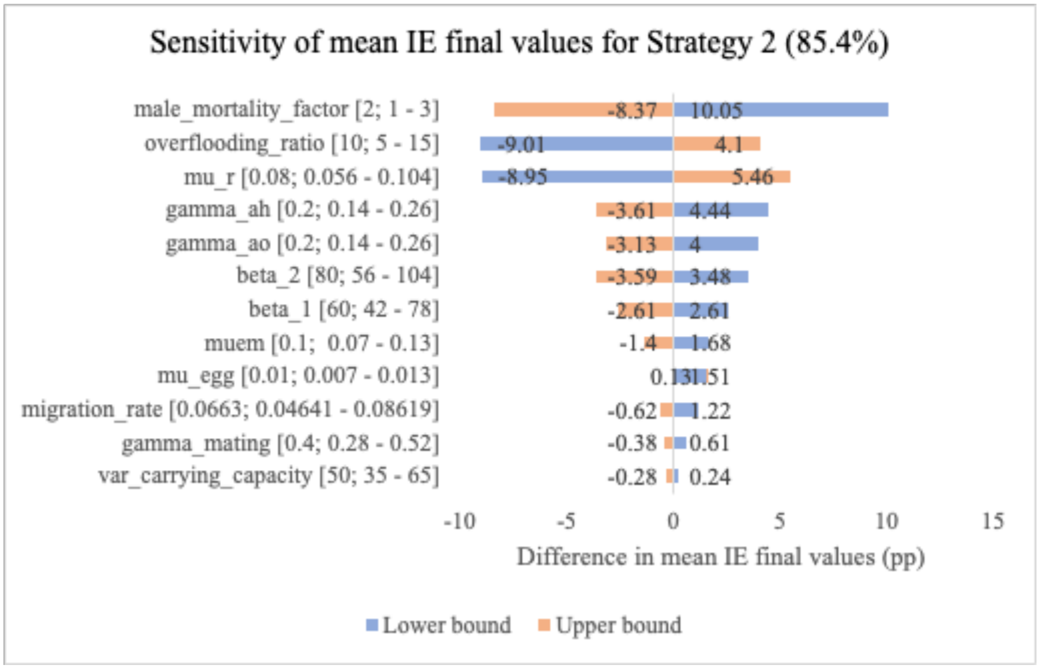
Sensitivity of mean IE final values for Strategy 2 (85.4%)
male_mortality_factor [2; 1 - 3]
overflooding_ratio [10; 5 - 15]
mu_r [0.08; 0.056 - 0.104]
gamma_ah [0.2; 0.14 - 0.26]
gamma_ao [0.2; 0.14 - 0.26]
beta_2 [80; 56 - 104]
beta_1 [60; 42 - 78]
muem [0.1; 0.07 - 0.13]
mu_egg [0.01; 0.007 - 0.013]
migration_rate [0.0663; 0.04641 - 0.08619]
gamma_mating [0.4; 0.28 - 0.52]
var_carrying_capacity [50; 35 - 65]
-8.37
10.05
-9.01
4.1
-8.95
5.46
-3.61
4.44
-3.13
4
-3.59
3.48
-2.61
2.61
-1.4
1.68
0.13
1.51
-0.62
1.22
-0.38
0.61
-0.28
0.24
-10
-5
0
5
10
15
Difference in mean IE final values (pp)
Lower bound
Upper bound

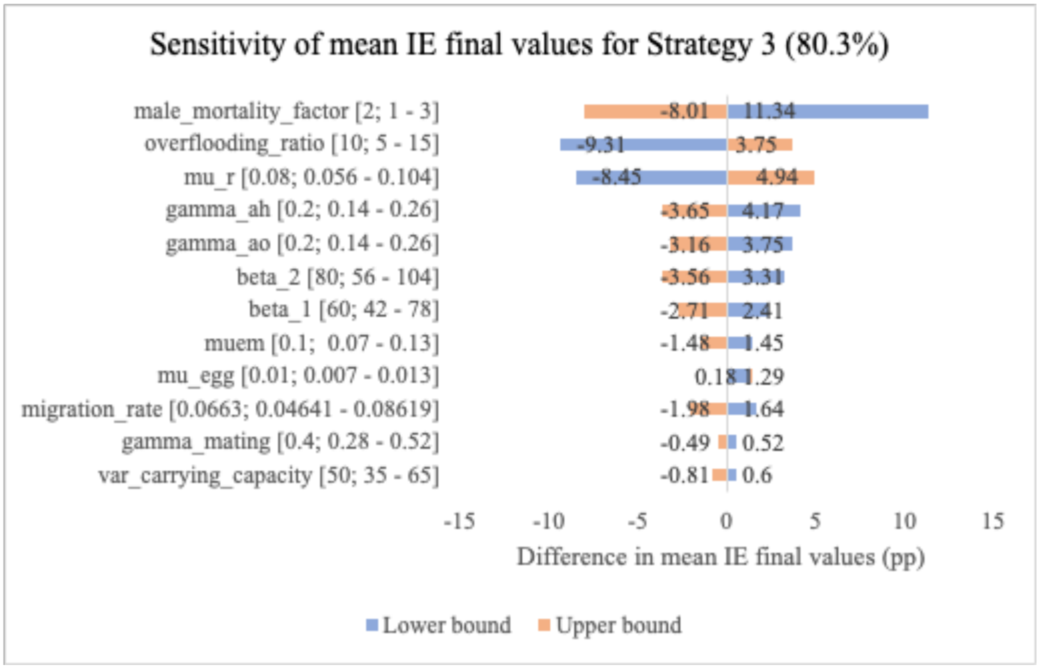
Sensitivity of mean IE final values for Strategy 3 (80.3%)
male_mortality_factor [2; 1 - 3]
overflooding_ratio [10; 5 - 15]
mu_r [0.08; 0.056 - 0.104]
gamma_ah [0.2; 0.14 - 0.26]
gamma_ao [0.2; 0.14 - 0.26]
beta_2 [80; 56 - 104]
beta_1 [60; 42 - 78]
muem [0.1; 0.07 - 0.13]
mu_egg [0.01; 0.007 - 0.013]
migration_rate [0.0663; 0.04641 - 0.08619]
gamma_mating [0.4; 0.28 - 0.52]
var_carrying_capacity [50; 35 - 65]
-8.01
11.34
-9.31
3.75
-8.45
4.94
-3.65
4.17
-3.16
3.75
-3.56
3.31
-2.71
2.41
-1.48
1.45
0.18
1.29
-1.98
1.64
-0.49
0.52
-0.81
0.6
-15
-10
-5
0
5
10
15
Difference in mean IE final values (pp)
Lower bound
Upper bound

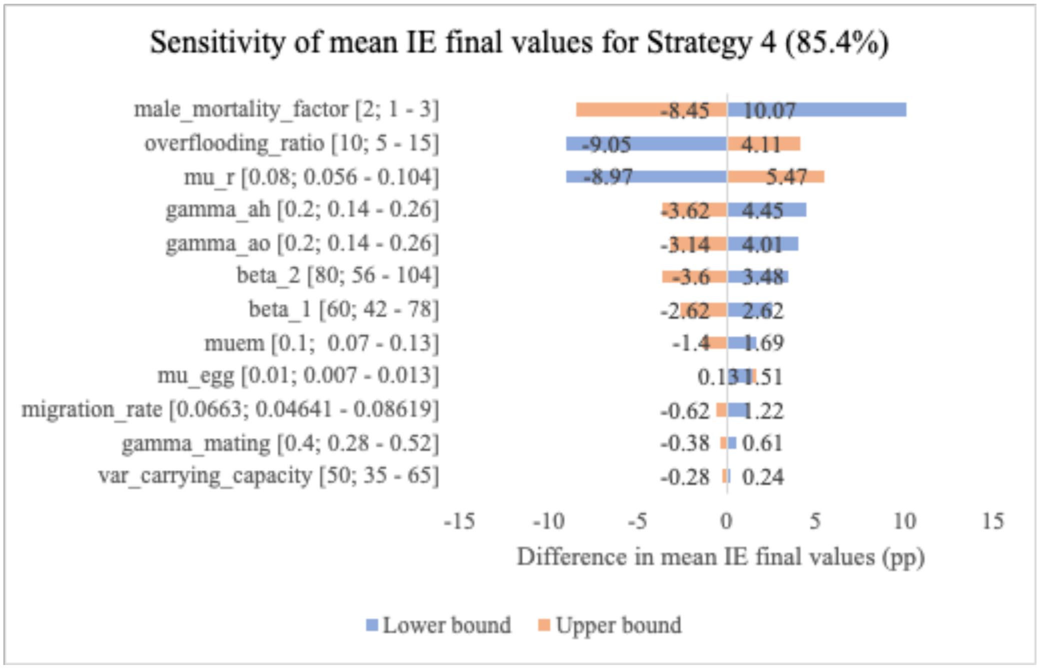
Sensitivity of mean IE final values for Strategy 4 (85.4%)
male_mortality_factor [2; 1 - 3]
overflooding_ratio [10; 5 - 15]
mu_r [0.08; 0.056 - 0.104]
gamma_ah [0.2; 0.14 - 0.26]
gamma_ao [0.2; 0.14 - 0.26]
beta_2 [80; 56 - 104]
beta_1 [60; 42 - 78]
muem [0.1; 0.07 - 0.13]
mu_egg [0.01; 0.007 - 0.013]
migration_rate [0.0663; 0.04641 - 0.08619]
gamma_mating [0.4; 0.28 - 0.52]
var_carrying_capacity [50; 35 - 65]
-8.45
10.07
-9.05
4.11
-8.97
5.47
-3.62
4.45
-3.14
4.01
-3.6
3.48
-2.62
2.62
-1.4
1.69
0.13
1.51
-0.62
1.22
-0.38
0.61
-0.28
0.24
-15
-10
-5
0
5
10
15
Difference in mean IE final values (pp)
Lower bound
Upper bound

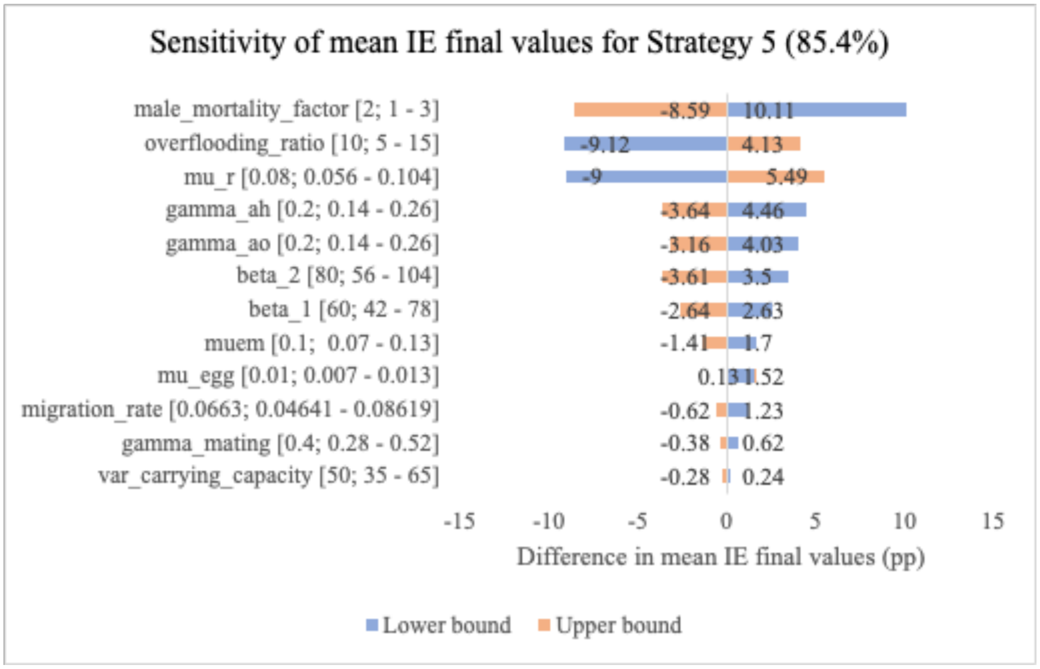
Sensitivity of mean IE final values for Strategy 5 (85.4%)
male_mortality_factor [2; 1 - 3]
overflooding_ratio [10; 5 - 15]
mu_r [0.08; 0.056 - 0.104]
gamma_ah [0.2; 0.14 - 0.26]
gamma_ao [0.2; 0.14 - 0.26]
beta_2 [80; 56 - 104]
beta_1 [60; 42 - 78]
muem [0.1; 0.07 - 0.13]
mu_egg [0.01; 0.007 - 0.013]
migration_rate [0.0663; 0.04641 - 0.08619]
gamma_mating [0.4; 0.28 - 0.52]
var_carrying_capacity [50; 35 - 65]
-8.59
10.11
-9.12
4.13
-9
5.49
-3.64
4.46
-3.16
4.03
-3.61
3.5
-2.64
2.63
-1.41
1.7
0.13
1.52
-0.62
1.23
-0.38
0.62
-0.28
0.24
-15
-10
-5
0
5
10
15
Difference in mean IE final values (pp)
Lower bound
Upper bound

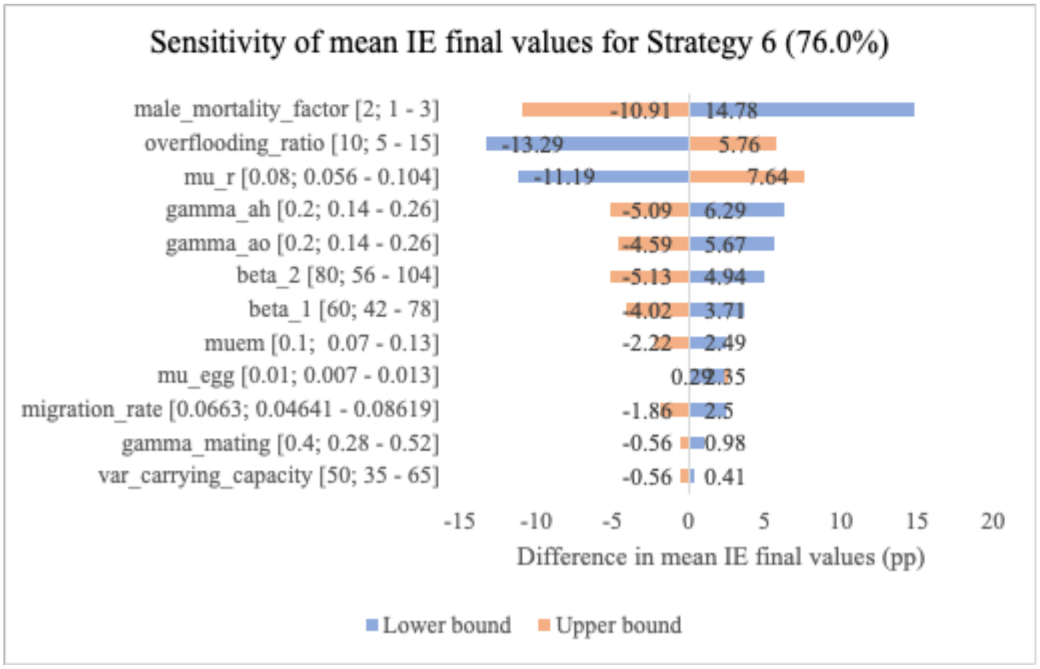
Sensitivity of mean IE final values for Strategy 6 (76.0%)
male_mortality_factor [2; 1 - 3]
overflooding_ratio [10; 5 - 15]
mu_r [0.08; 0.056 - 0.104]
gamma_ah [0.2; 0.14 - 0.26]
gamma_ao [0.2; 0.14 - 0.26]
beta_2 [80; 56 - 104]
beta_1 [60; 42 - 78]
muem [0.1; 0.07 - 0.13]
mu_egg [0.01; 0.007 - 0.013]
migration_rate [0.0663; 0.04641 - 0.08619]
gamma_mating [0.4; 0.28 - 0.52]
var_carrying_capacity [50; 35 - 65]
-10.91
14.78
-13.29
5.76
-11.19
7.64
-5.09
6.29
-4.59
5.67
-5.13
4.94
-4.02
3.71
-2.22
2.49
0.29
2.35
-1.86
2.5
-0.56
0.98
-0.56
0.41
-15
-10
-5
0
5
10
15
20
Difference in mean IE final values (pp)
Lower bound
Upper bound

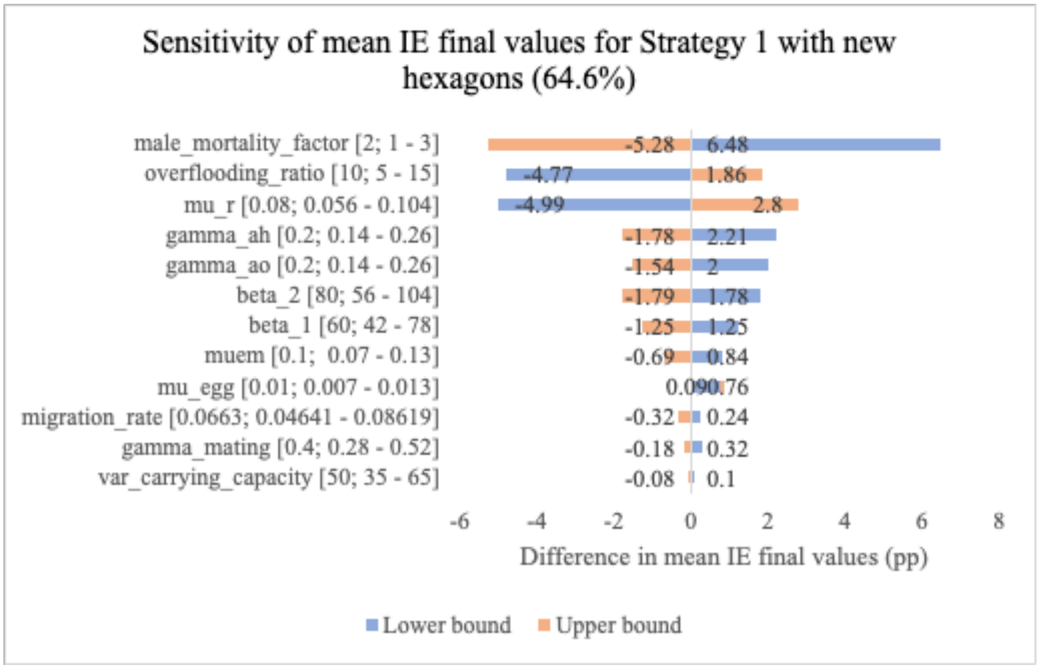
Sensitivity of mean IE final values for Strategy 1 with new hexagons (64.6%)
male_mortality_factor [2; 1 - 3]
overflooding_ratio [10; 5 - 15]
mu_r [0.08; 0.056 - 0.104]
gamma_ah [0.2; 0.14 - 0.26]
gamma_ao [0.2; 0.14 - 0.26]
beta_2 [80; 56 - 104]
beta_1 [60; 42 - 78]
muem [0.1; 0.07 - 0.13]
mu_egg [0.01; 0.007 - 0.013]
migration_rate [0.0663; 0.04641 - 0.08619]
gamma_mating [0.4; 0.28 - 0.52]
var_carrying_capacity [50; 35 - 65]
-5.28
6.48
-4.77
1.86
-4.99
2.8
-1.78
2.21
-1.54
2
-1.79
1.78
-1.25
1.25
-0.69
0.84
-0.32
0.24
-0.18
0.32
-0.08
0.1
Difference in mean IE final values (pp)
Lower bound
Upper bound

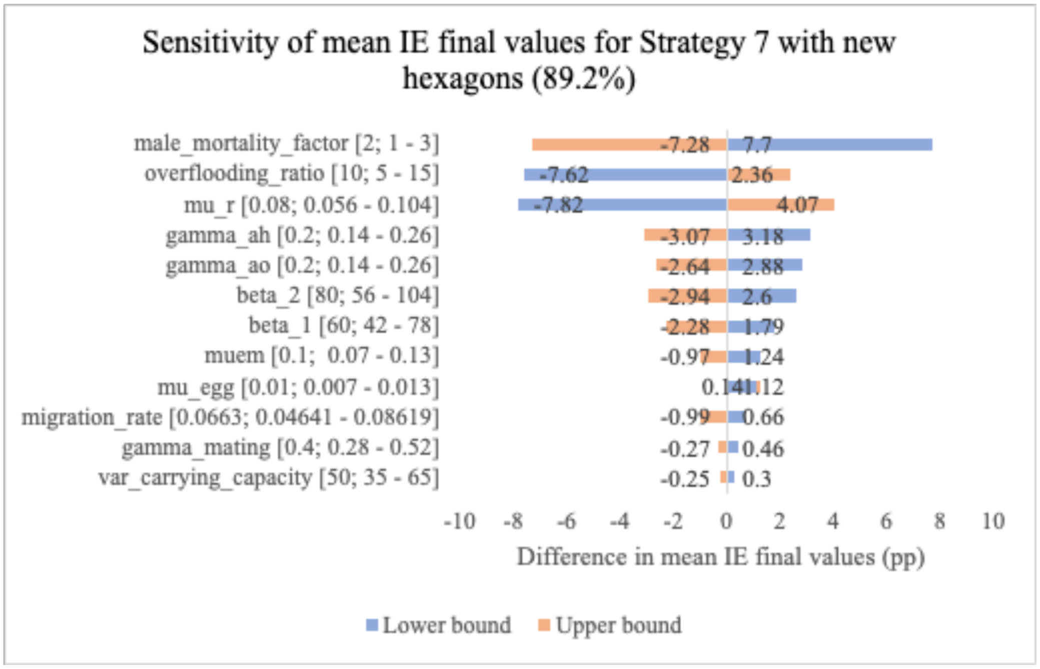
Sensitivity of mean IE final values for Strategy 7 with new hexagons (89.2%)
male_mortality_factor [2; 1 - 3]
overflooding_ratio [10; 5 - 15]
mu_r [0.08; 0.056 - 0.104]
gamma_ah [0.2; 0.14 - 0.26]
gamma_ao [0.2; 0.14 - 0.26]
beta_2 [80; 56 - 104]
beta_1 [60; 42 - 78]
muem [0.1; 0.07 - 0.13]
mu_egg [0.01; 0.007 - 0.013]
migration_rate [0.0663; 0.04641 - 0.08619]
gamma_mating [0.4; 0.28 - 0.52]
var_carrying_capacity [50; 35 - 65]
-7.28
7.7
-7.62
2.36
-7.82
4.07
-3.07
3.18
-2.64
2.88
-2.94
2.6
-2.28
1.79
-0.97
1.24
-0.99
0.66
-0.27
0.46
-0.25
0.3
Difference in mean IE final values (pp)
Lower bound
Upper bound

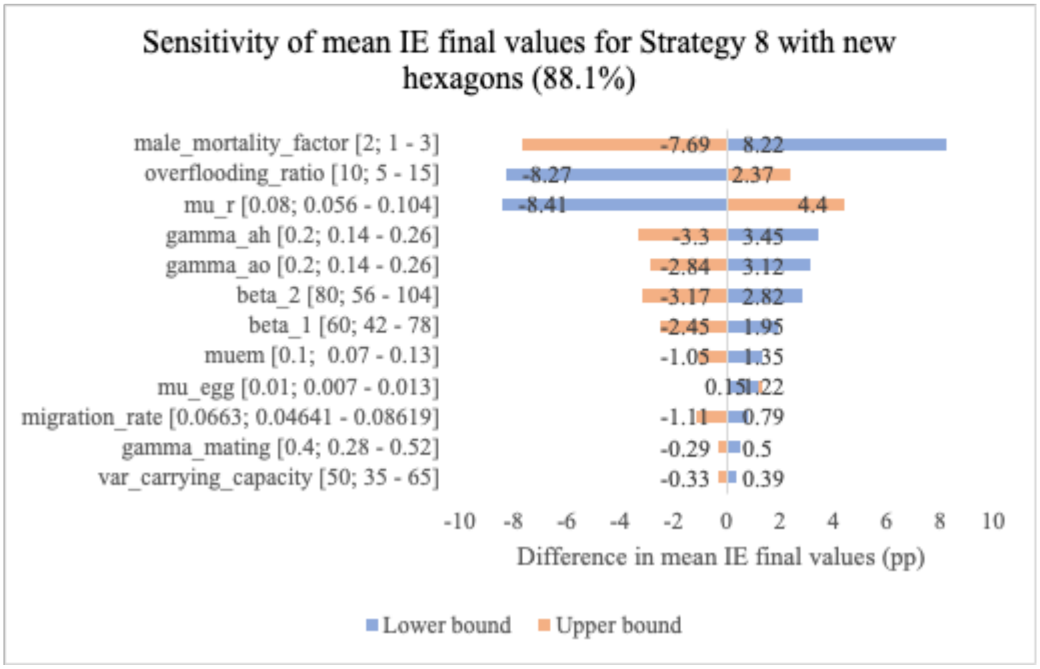

Sensitivity of mean IE final values for Strategy 8 with new hexagons (88.1%)
male_mortality_factor [2; 1 - 3]
overflooding_ratio [10; 5 - 15]
mu_r [0.08; 0.056 - 0.104]
gamma_ah [0.2; 0.14 - 0.26]
gamma_ao [0.2; 0.14 - 0.26]
beta_2 [80; 56 - 104]
beta_1 [60; 42 - 78]
muem [0.1; 0.07 - 0.13]
mu_egg [0.01; 0.007 - 0.013]
migration_rate [0.0663; 0.04641 - 0.08619]
gamma_mating [0.4; 0.28 - 0.52]
var_carrying_capacity [50; 35 - 65]
-7.69
8.22
-8.27
2.37
-8.41
4.4
-3.3
3.45
-2.84
3.12
-3.17
2.82
-2.45
1.95
-1.05
1.35
0.15
1.22
-1.11
0.79
-0.29
0.5
-0.33
0.39
-10 -8 -6 -4 -2 0 2 4 6 8 10
Difference in mean IE final values (pp)
Lower bound
Upper bound

**Table S3. List of subzones in the model validation study.**
Model validation subzones have a total population of 566,990.

| Zone | Subzone | Population size |
|---|---|---|
| Tampines | Tampines North | 25680 |
| Tampines | Tampines East | 126460 |
| Tampines | Tampines West | 81370 |
| Yishun | Yishun West | 52160 |
| Yishun | Yishun Central | 2960 |
| Yishun | Yishun South | 41920 |
| Yishun | Yishun East | 70190 |
| Yishun | Northland | 27280 |
| Choa Chu Kang | Yew Tee | 40070 |
| Choa Chu Kang | Peng Siang | 34250 |
| Choa Chu Kang | Keat Hong | 38990 |
| Bukit Batok | Hong Kah North | 25660 |

**Table S4. List of subzones in the scale-down strategies study.**
Current release subzones have a total population of 1,304,700.

| Zone | Subzone | Population size |
|---|---|---|
| Tampines | Tampines North | 25680 |
| Tampines | Tampines East | 126460 |
| Tampines | Tampines West | 81370 |
| Woodlands | Midview | 34580 |
| Woodlands | Woodlands East | 100220 |
| Yishun | Yishun West | 52160 |
| Yishun | Yishun Central | 2960 |
| Yishun | Yishun South | 41920 |
| Yishun | Yishun East | 70190 |
| Yishun | Northland | 27280 |
| Sengkang | Sengkang Town Centre | 61210 |
| Sengkang | Rivervale | 59180 |
| Marine Parade - Mountbatten | Mountbatten | 10260 |
| Marine Parade - Mountbatten | Katong | 9450 |
| Marine Parade - Mountbatten | Frankel | 35010 |

| | | |
|---|---|---|
| Punggol | Punggol Field | 49010 |
| Hougang | Hougang West | 44840 |
| Holland | Ulu Pandan | 11470 |
| Geylang - MacPherson | MacPherson | 27120 |
| Commonwealth | Holland Drive | 12210 |
| Commonwealth | Mei Chin | 16590 |
| Clementi - West Coast | West Coast | 10100 |
| Clementi - West Coast | Clementi Central | 14780 |
| Clementi - West Coast | Clementi North | 29400 |
| Choa Chu Kang | Yew Tee | 40070 |
| Choa Chu Kang | Peng Siang | 34250 |
| Choa Chu Kang | Keat Hong | 38990 |
| Bukit Merah - Telok Blangah | Alexandra Hill | 13220 |
| Bukit Merah - Telok Blangah | Redhill | 11090 |
| Bukit Merah - Telok Blangah | Henderson Hill | 13000 |
| Bukit Merah - Telok Blangah | Tiong Bahru Station | 14870 |
| Bukit Merah - Telok Blangah | Tiong Bahru | 12240 |
| Bukit Merah - Telok Blangah | Kampong Tiong Bahru | 8840 |
| Bukit Merah - Telok Blangah | Telok Blangah Rise | 11780 |
| Bukit Merah - Telok Blangah | Telok Blangah Way | 8590 |
| Bukit Batok | Hong Kah North | 25660 |
| Bedok | Kaki Bukit | 35880 |
| Bedok | Bedok North | 82770 |

**Table S5. List of new subzones in the redistribution strategies study.**
New release subzones have a total population of 615,130 people.

| **Zone** | **Subzone** | **Population size** |
|---|---|---|
| Jurong West | Yunnan | 66890 |
| Jurong West | Jurong West Central | 62960 |
| Jurong West | Hong Kah | 53380 |
| Jurong West | Boon Lay Place | 28590 |
| Jurong West | Taman Jurong | 39600 |
| Jurong West | Wenya | 8280 |

| | | |
|---|---|---|
| Toa Payoh | Toa Payoh Central | 27570 |
| Novena | Balestier | 33040 |
| Kallang | Bendemeer | 37420 |
| Ang Mo Kio | Cheng San | 27280 |
| Ang Mo Kio | Chong Boon | 25680 |
| Ang Mo Kio | Yio Chu Kang West | 23820 |
| Ang Mo Kio | Kebun Bahru | 22250 |
| Ang Mo Kio | Townsville | 20210 |
| Ang Mo Kio | Shangri-La | 17630 |
| Pasir Ris | Pasir Ris West | 35580 |
| Pasir Ris | Pasir Ris Drive | 52650 |
| Pasir Ris | Pasir Ris Central | 32300 |

### Parameter estimation

#### Estimation of the relative ratio between fixed and variable carrying capacities

We assumed the relative ratio between human- to precipitation- dependent carrying capacities to be 2:1 based on a Singapore National Environmental Agency report of the number of mosquito breeding sites in residential and non-residential areas (*Mosquito Breeding Habitats Found in Homes Doubled in January Compared to Last Year*, 2024). Specifically, we assumed that breeding sites found in homes were human-dependent and those not in homes were precipitation-dependent.

#### Estimation of the adult male mortality rate process-based function

Magori et al. (2009) assumed that the adult male mortality probability was about twice that of adult females. Therefore, we assumed an adult male mortality rate with a two-time multiplier compared to the adult female mortality rate function.

#### Estimation of migration rate, $\upsilon$

The Singapore National Environmental Agency reported that almost all male IIT mosquitoes were caught within 40 meters from where they were released, and that the mosquitoes had a median post-release lifespan of 4 days (*Phase 1 Small-Scale Field Study*, n.d.). From this, we assumed that the mean horizontal travel distance for all IIT mosquitoes was 40 meters, and determined that the mean lifespan was 5.77 days by using the following formula, assuming that mosquito longevity followed an exponential decay:

$$\text{Mean lifespan} = \text{half life} / \log(2)$$

We then derived the mean daily horizontal travel distance by assuming that mosquito dispersal followed a 2D Gaussian distribution, where the average distance from the mean over 5.77 days was 40 meters. The single-day horizontal travel distance of 16.65 meters was derived as such:

$$\text{40 meters} = \sqrt{5.77({\sigma_x}^2 + {\sigma_y}^2)}$$

$$\text{Daily average distance} = \sqrt{{\sigma_x}^2 + {\sigma_y}^2}$$

$$= 16.65 \text{ meters}$$

The probability of a mosquito leaving the hexagon was determined by running a simulation with 1000 iterations, where in each iteration, a random point within the hexagon corresponding to the initial mosquito position and a random direction representing the single-day direction of travel was selected. The single-day probability that a mosquito would leave a hexagon was the proportion of iterations where the final mosquito position was outside of the bounds of the hexagon. This simulation was repeated 1000 times, and a mean value of 6.63% was obtained.
We assumed this migration rate applied to all adult mosquitoes in this study.

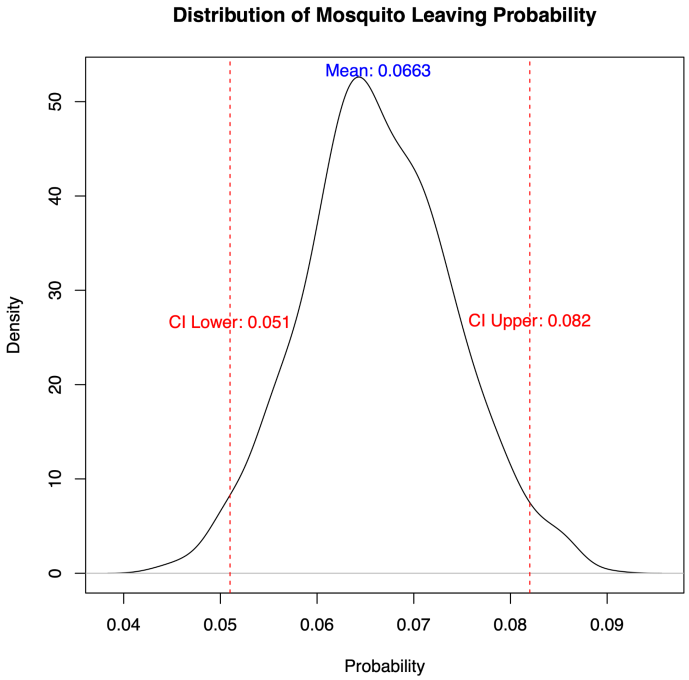


**Figure S1. Distribution of the probability of mosquitoes leaving a hexagon.**
